# A conserved electron transport chain sensitizes Bacteroides to a pore-forming type VI secretion toxin

**DOI:** 10.1101/2025.08.29.672963

**Authors:** Hannah K. Ratner, Brandon D. Duong, Pengrui Miao, Savannah K. Bertolli, Beth A. Shen, Uma Mitchell, Larry A Gallagher, Matthew Radey, S Brook Peterson, Joseph D. Mougous

## Abstract

Data suggest that antagonism between bacteria is prevalent within the gut microbiome. Such antagonism could have profound consequences on the fitness of species; however, the susceptibility determinants to even the most pervasive antagonistic factors in this ecosystem remain incompletely understood. Here, we screened for genetic factors that impact the susceptibility of *Bacteroides* to type VI secretion system (T6SS)-delivered toxins. This revealed that the Bte2 family of pore-forming toxins, which are widespread in *B. fragilis* and other human gut-associated Bacteroidales, strictly require the H^+^/Na^+^-translocating ferredoxin:NAD^+^ reductase (Rnf) electron transport chain within target cells in order to intoxicate. In *Bacteriodes*, the precise metabolic role of the conserved Rnf pathway has not been defined. We establish that the Rnf complex is important for redox balancing within cells utilizing sugars derived from dietary fiber and is critical for fitness in vivo. Surprisingly, we find that while the intact Rnf membrane complex is required for Bte2 intoxication, Rnf-catalyzed electron transport is dispensable. We propose that the Rnf complex facilitates Bte2 membrane insertion, leading to intoxication via membrane depolarization. Our data suggest that T6SS toxins may avoid collateral damage within a complex ecosystem by recognizing discriminatory features of competitor species.

**Significance:** Pathways for interbacterial antagonism are prevalent in the gut microbiome. The breadth of targeting and specificity determinants of these systems remain largely uncharacterized. We discovered that a widespread pore-forming toxin produced by gut Bacteroidales requires the conserved Rnf protein complex in target organisms. Although this complex is dispensable during *in vitro* growth, we show it is required for *Bacteroides* fitness during colonization of the mammalian gut. Our data support a model in which transient interaction between the toxin and Rnf enables rearrangement of the protein, facilitating membrane insertion. Related toxins found in Proteobacteria lack the requirement for Rnf, suggesting that competition between Bacteroidales species in the gut may be driving specialization of their antibacterial toxins.

## Introduction

Competition among bacteria is an ancient and ubiquitous process that has selected for a fascinating variety of antagonistic mechanisms (1). Among these is the Gram-negative bacterial type VI secretion system (T6SS), a multi-protein assemblage that is capable of catalyzing the contact-dependent transfer of toxic effector proteins into neighboring bacteria (2). Acting via a contractile phage-like mechanism, the T6SS breaches the outer-membrane of target Gram-negative cells, releasing a cocktail of effectors into the periplasm (3, 4). The toxins subsequently act within this subcellular compartment or translocate across the cytoplasmic membrane to reach their substrates (2). By functioning in a receptor-independent manner, and by delivering a multitude of toxins concurrently, the T6SS is able to both target a broad range of organisms and avoid the rapid evolution of resistance (5-7).

The unique features and fitness benefits afforded by the T6SS are reflected in its broad distribution across Gram-negative phyla, including the phyum Bacteroidota (8). Members of this phylum, in particular those within the order Bacteroidales, are adapted to the mammalian digestive tract. Indeed, Bacteroidales constitute 20-80% of all bacteria in gut microbiome of a healthy human(9). Though its role in interbacterial antagonism via toxin delivery is conserved, the Bacteroidales T6SS (subtype 3, T6SS^iii^) is distinct from the canonical systems first described in Proteobacteria (subtype 1, T6SS^i^). For example, the Bacteroidales T6 apparatus lacks orthologs of three strictly conserved proteins constituting the Proteobacterial T6SS membrane complex, and instead possesses five unrelated proteins that form a functionally equivalent structure (8, 10). Also, whereas Proteobacterial T6SSs are typically components of the core genome, those in the Bacteroidales are most often located on mobile elements (11). An exception are the genetic architecture 3 (GA3) T6SSs restricted to *Bacteroides fragilis* (7). A number of the differences between the systems of these organisms relate to the toxins they deliver. Many of the those delivered by the Bacteroidales T6SS lack homologs outside of the order and, correspondingly, appear to intoxicate via distinctive mechanisms (11-13). A recent study found that one such toxin, Bte1 from *B. fragilis* NCTC9343, interacts with a periplasmic chaperone complex within target cells, leading to a proposed mechanism of toxicity involving widespread misfolding of nascent cell envelope proteins (14). The unique adaptations of the Bacteroidales T6SS are likely to reflect parameters specific to the environmental conditions and bacterial community composition within the mammalian gut microbiome.

Pore formation is a prevalent mechanism of action by interbacterial toxins, and several evolutionary distinct T6SS pore forming toxin families are currently recognized (6, 15-17). Among these are the VasX family toxins which depolarize the inner-membrane of target cells by introducing small, ion-selective pores. Though the mechanism by which these large multi-domain proteins introduce pores remains unknown, recent structures of VasX toxins Ptx2 and Tke5, from *Pseudomonas aeruginosa* and *P. putida*, respectively, suggest that membrane sensing following periplasmic delivery triggers deployment of the transmembrane domains (18, 19). Here we show that the Bacteroidales T6SS effector Bte2 is a divergent member of the VasX toxin family that requires the conserved membrane components of the Rnf electron transport chain within recipient cells in order to intoxicate. We establish a critical role for Rnf in Bacteroidales during growth on sugars derived from dietary fibers and provide bioinformatic evidence that Bte2-dependence on Rnf is limited to gut Bacteroidales. Our findings suggest that in contrast to the broad, receptor-independent targeting observed for the T6SS^i^, the Bacteroidales T6SS^iii^ is specifically adapted to mediate competition with other *Bacteroides* species, their most frequent competitors in the gut.

## Results

### The Rnf Pathway sensitizes *Bacteroides* species to a family of T6SS toxins

To identify genes that mediate susceptibility to T6SS toxins in *Bacteroides* species, we performed random barcode transposon site sequencing (RB-Tn-seq) screens using *Bacteroides thetaiotaomicron* (*B. theta*) as the target (20, 21). We utilized two *B. fragilis* strains that possess different T6SS toxin repertoires in these experiments: ATCC NCTC9343, which encodes the periplasmic chaperone inhibitor Bte1 and a second toxin of unknown function (Bte2^9343^), and 638R, which encodes an ortholog bearing 30% amino acid identity to Bte2^9343^, referred to hereafter as Bte2^638R^, and an additional toxin of unknown function, which we rename here Bte3 (8, 22). In these screens, the *B. theta* transposon mutant library was grown under T6SS-conducive conditions (cell-cell contact promoting) in co-culture with wild-type *B. fragilis*, strains engineered to deliver individual toxins, or T6SS-deficient control strains. We then identified *B. theta* genes in which transposon mutants were selected during antagonism, which represent candidate toxin susceptibility determinants (Table S1).

Exposure to distinct *B. fragilis* toxins resulted in the enrichment of mutations in different genes (Fig. 1A-D, S1A-D, Table S1). Mutations in a limited number of genes were selected by more than one toxin, suggesting they encode proteins that play a general role in sensitizing *B. theta* to T6SS-mediated intoxication. Supporting this, mutations in a subset of these genes were also selected during competition with the wild-type *B. fragilis* strains, which deliver both of their toxins (Fig. S1F-G, Table S1). Mutations selected most strongly by antagonism by both wild-type *B. fragilis* strains resided within genes encoding a predicted AAA+ family ATPase (BT_RS24785), a hypothetical protein (BT_RS07405), and a glycosyltransferase family 2 protein (BT_RS17040) (Fig. S1F-G). Genes in which mutations were specifically selected during intoxication by Bte1 encode largely uncharacterized proteins (Fig. 1A). Notably, mutations in the gene encoding the periplasmic Bte1 interaction partner, *ppiD*, were not enriched, consistent with the strong growth defect observed for PpiD inactivation (14). In screens with Bte3, the gene with the strongest enrichment of mutations was *batD*, which is present in an operon implicated in aerotolerance (Fig. 1B). Other genes in this locus exhibited low insertion density, consistent with their general importance for *Bacteroides* species physiology (Table S1, (23)). The mechanism by which the products of this operon promote *Bacteroides* species survival during oxygen exposure is not known, but similar operons have been linked to stress resistance in other species (24, 25).

**Figure 1.**
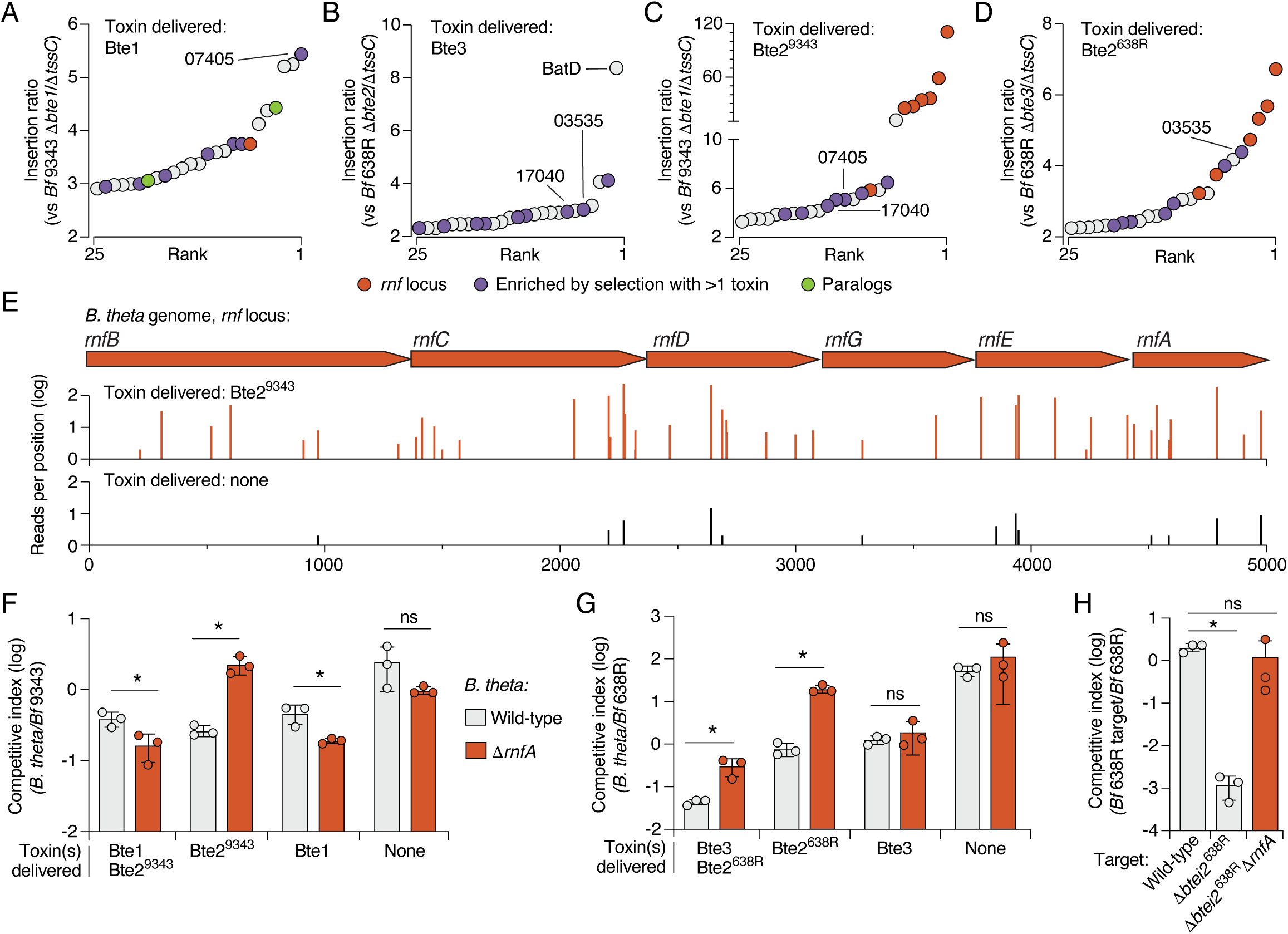
The Rnf complex sensitizes *Bacteroides* spp. to a family of *B. fragilis* toxins. **A-D)** Rank fold enrichment of *B. theta* (*Bt*) transposon insertion mutants following growth in competition with *B. fragilis* delivering the indicated T6SS toxins. Mutants specifically mentioned in the text are highlighted. BT_RS07405, hypothetical; BT_RS17040, predicted glycosyltransferase; BT_RS03535, N-acetylmuramoyl-L-alanine amidase. **E)** Schematic of genes with the Rnf locus indicating transposon insertion sites and normalized insertion frequency following exposure to Bte2^9343^ (top) or no toxins (bottom). **F-G)** Outcome of growth competition assays between *Bt* (wild-type or Δ*rnfA*) and the indicated strains of *B. fragilis* 9343 (F) or 638R (G). Data represent mean ± SD (*p<0.05, unpaired two-tailed Student’s *t*-test (n=3).

Screens with both Bte2 family members (Bte2^638R^ and Bte2^9343^) revealed striking enrichment of mutations in genes encoding components of the H⁺/Na⁺-translocating ferredoxin:NAD⁺ reductase (Rnf) complex (Fig. 1C-E, S1A-C). Mutations within all genes of the Rnf locus were enriched during exposure to one or both Bte2 orthologs, and constituted those most enriched by Bte2 intoxication by either strain. To validate this finding, we inactivated Rnf in *B. theta* via in-frame deletion of *rnfA* and performed pairwise growth competitions with *B. fragilis* strains delivering each toxin. Consistent with the results of our Tn-seq screens, *B. theta* Δ*rnfA* exhibits increased fitness during competitive growth assays with *B. fragilis* producing Bte2^638R^ or Bte2^9343^, but not Bte1 or Bte3 (Fig. 1F-G, S1B-D). Indeed, in both our Tn-seq and pairwise experiments, *B. theta* strains lacking Rnf functionality exhibited a modest fitness deficit relative to the wild-type when challenged with Bte1. To determine whether the Rnf dependence observed for Bte2 family toxins extends to more distantly related orthologs, we measured the impact of Rnf-inactivation on *B. theta* competitiveness with *B. fragilis* DS-71, which encodes a Bte2 ortholog from a distinct clade (Fig. S1G). Rnf inactivation similarly yielded a fitness benefit to *B. theta* in competitive growth assays with this strain (Fig. S1H).

Given that the Rnf complex is broadly conserved across Bacteroidales, we next asked whether it is generally required for Bte2-mediated intoxication. We generated an in-frame deletion of *rnfA* in a *B. fragilis* 638R derivative sensitized to Bte2^638R^ via deletion of the genes encoding the toxin and its adjacent immunity determinant (Δ*bte/i*2). In competition with the parent strain, *B. fragilis* Δ*bte/i2* displayed a pronounced competitive defect (Fig. 1H). However, inactivation of *rnfA* in this strain granted full protection from Bte2-based intoxication (Fig. 1H) Together, our data suggest that intoxication by distantly related Bte2 family members requires the presence of the Rnf pathway in multiple target species.

### The Rnf complex is required for *Bacteroides* survival in the mammalian gut

Toxins mediating interbacterial antagonism typically subvert the emergence of resistance by targeting essential molecules. Thus, we were surprised to discover that inactivation of *rnf* genes, which apparently does not confer a fitness cost during standard *in vitro* propagation, leads to Bte2 resistance. Given the widespread conservation of the Rnf complex among *Bacteroidales* and the abundance of genes encoding Bte2 family members in gut microbiome gene catalogs, we hypothesized that the physiological importance of the Rnf complex for *Bacteroidales* growth in their native environment may not be captured by our *in vitro* growth conditions.

Although the function of the Rnf complex in *Bacteroidales* is unknown, biochemical analyses in *B. fragilis* and metabolic predictions and metabolite measurements in related gut anaerobes in the *Prevotella* genus suggest that it serves to couple sodium extrusion to the transfer of electrons from reduced ferredoxin to NAD^+^, thereby contributing to the chemiosmotic gradient and regenerating cofactors required for fermentative growth (Fig. 2A) (26, 27). *B. theta* is a metabolically versatile species able to use a wide range of growth substrates (28-31). Given that catabolic pathways can vary in their requirements for NADH and reduced ferredoxin, we assessed the importance of the Rnf complex for *B. theta* growth on different carbon sources representative of substrates it likely encounters in the human gut. These included common dietary sugars (glucose, fructose) and monosaccharides derived from dietary fiber (galacturonic acid, rhamnose). As in rich media, inactivation of Rnf did not significantly impact *B. theta* growth in defined minimal media (DMM) containing sugars commonly consumed as monosaccharides (Fig. 2B). However, we observed a pronounced growth defect of *B. theta* 11*rnfA* in DMM containing galacturonic acid and rhamnose. During examination of the predicted metabolic pathways for utilization of these substrates, we noted that early steps in galacturonic acid and rhamnose catabolism require NADH, and that there is not a downstream conversion that regenerates the electron donor (Figure S2A). Rnf could thus be important for growth on these substrates by using electrons from reduced ferredoxin to replenish the NADH pool, similar to the role it plays in redox balancing during nitrogen fixation (32, 33).

1. *B. theta* primarily employs mixed acid fermentation, which generates a range of secreted products, including acetate, formate, succinate, propionate, and smaller amounts of lactate and certain amino acids (34-36). Previous studies have shown that high exogenous concentrations of fermentation products can exhibit feedback inhibition on certain metabolic pathways in *B. theta,* thus altering the cellular redox requirements (37). Accordingly, we examined whether the addition of fermentation products to DMM alters the requirement for Rnf function in *B. theta*. During growth on glucose, we observed a modest reduction in growth yield for *B. theta* 11*rnf* upon propionate supplementation (Fig. 2C). We speculate that excess propionate exhibits feedback inhibition on one or more step in the succinate pathway, resulting in increased flux toward acetate production from pyruvate (Figure S2). This pathway requires pyruvate:ferredoxin oxidoreductase, which generates reduced ferredoxin. In the absence of reoxidation by Rnf, accumulation of this molecule may retard growth. If the modest growth defects we observed for Rnf-inactive strains during supplementation with propionate or growth on galacturonic acid or rhamnose are due to redox imbalance, then we predicted that the combination of propionate supplementation and use of NADH-requiring growth substrates should exacerbate the fitness consequences of Rnf inactivation. Indeed, we find that during growth on galacturonic acid or rhamnose, supplementation with propionate resulted in acute toxicity (Fig. 2D). On DMM supplemented with galacturonic acid and proprionate, *B. theta* populations dropped to nearly undetectable levels; this medium is thus selective for Rnf-function, a feature we exploit below. This phenotype was complemented by expression of the locus from a neutral site on the chromosome (Figure 2D). Our *in vitro* growth assays suggest that in the mammalian gut, where dietary fibers are a common growth substrate and propionate can often be detected at high levels, the Rnf complex is crucial for *Bacteroides* species fitness (38-41).

**Figure 2.**
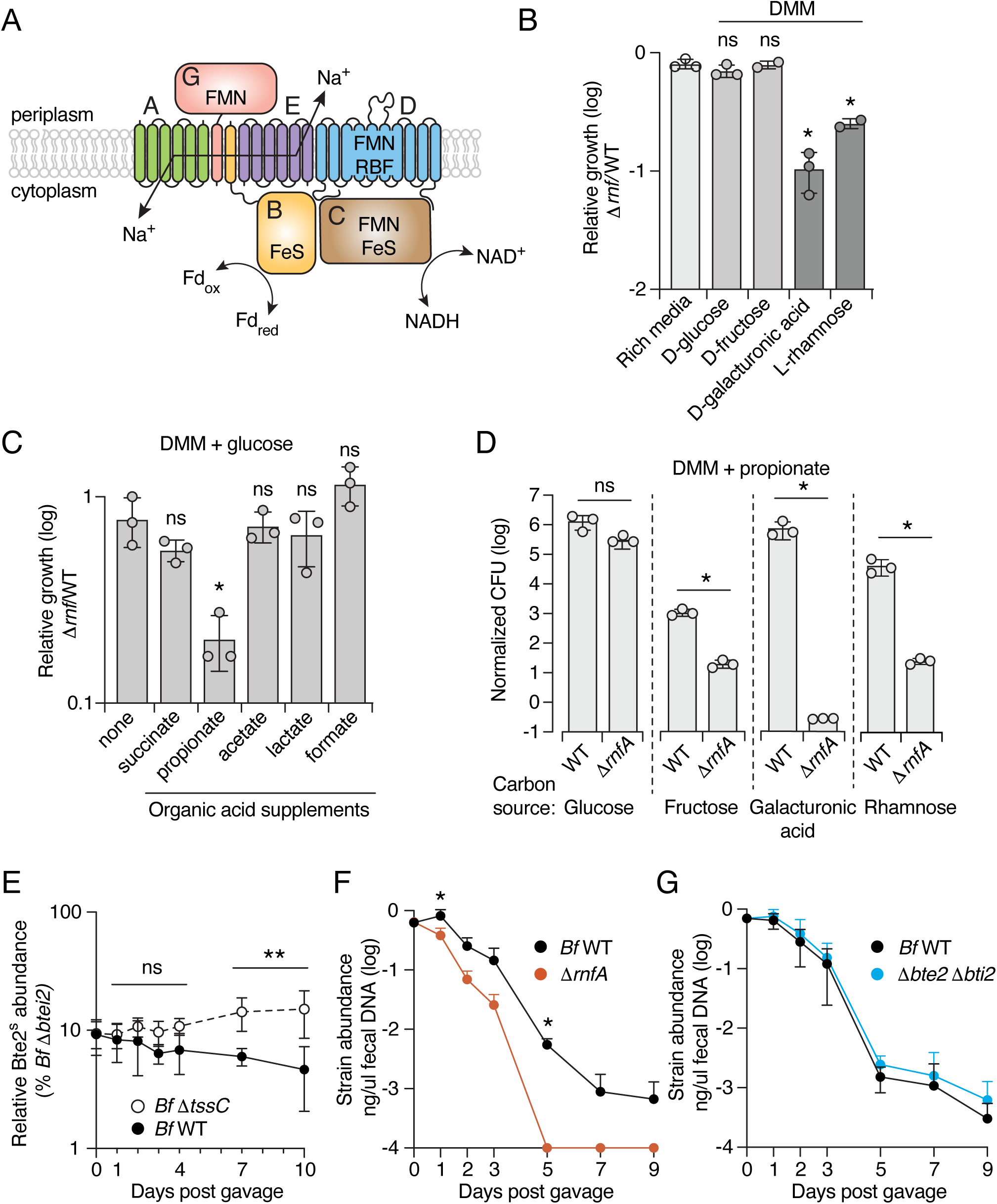
The Rnf complex enables *Bacteroides* to survive in the chemical environment of the mammalian gut. **A**) Schematic of the *B. theta* Rnf complex in the inner membrane, indicating the likely flow. **B-C**) Ratio of growth yields of *B. theta* Δ*rnfBCDGEA* compared to wild-type on DMM with different carbon sources (B) or DMM with glucose supplemented with the indicated fermentation end products (C). **D**) Normalized growth yields of *B. theta* wild-type or Δ*rnfBCDGEA* on DMM with propionate and the indicated monosaccharides. Data in B-D represent mean ± SD (*p<0.05, unpaired two-tailed Student’s *t*-test, n=3). **E**) Relative abundance of *B. fragilis* 11*bte2 bti2* (Bte2-sensitive) in mouse fecal pellets at the indicated time points post gavage with either *B. fragilis* 638R wild type or 11*tssC*. **F, G**) Abundance (ng/µl fecal DNA) of the indicated strains post co-gavage into gnotobiotic mice. Mice were simultaneously gavaged with a complex microbiota derived from wild mice (42). (**P<0.002, mean +/-SD, 2-way ANOVA, combined results from 2 biological replicates with n=5 per group per replicate)

For the Rnf complex to be retained by *Bacteroides* in the face of Bte2 antagonism in natural communities, we reasoned that the contribution of the complex to fitness must exceed the cost of Bte2 antagonism in this environment. To investigate the relative fitness consequences of Rnf-inactivation and Bte2-senisitization in a more physiologically relevant context than in vitro assays, we employed gnotobiotic mice fed a standard plant polysaccharide-rich chow diet and colonized with a complex microbiota derived from wild mice, WildR (42). We introduced this community via oral gavage, along with wild-type *B. fragilis* and either *B. fragilis* Δ*rnfA* or Δ*btei2* (1:1 initial ratio). We found *B. fragilis* Δ*rnfA* was at a lower abundance than wild-type in fecal samples after just one day post-gavage, and by day five, it was undetectable in the samples (Fig. 2F). In contrast, *B. fragilis* Δ*btei2* was maintained at similar levels to the wild-type over the duration of the experiment (Fig 2G).

One potential explanation for the lack of a fitness cost associated with sensitizing *B. fragilis* to Bte2 in our *in vivo* experiment could be that Bte2 is not active in this environment. However, it is well documented that T6SS-mediated antagonism is dampened in the context of a complex community and is likely relevant over longer timescales physiologically (22, 43-45). To more sensitively assess whether Bte2-mediated antagonism occurs between *B. fragilis* strains *in vivo*, we co-colonized gnotobiotic mice with wild-type *B. fragilis* 638R and our isogenic mutant sensitized to Bte2 (*B. fragilis* Δ*btei2*) at a 100:1 initial ratio in the absencZAe of the WildR community. As a control, we performed a similar experiment using a T6SS-inactive *B. fragilis* strain. After ten days of co-colonization, we found a significant fitness cost to Bte2 sensitization in the presence of wild-type, but not T6SS-inactive *B. fragilis* (Fig. 2E). This experiment demonstrates that Bte2-mediated antagonism between *Bacteroides* strains occurs *in vivo,* consistent with the *in vivo* competitive defect of a *B. fragilis* strain sensitized to both Bte1 and Bte2 previously observed (45). Overall, our *in vivo* experiments suggest that in their native environment, *Bacteroides* species utilize the Rnf complex for important metabolic transformations, and that the fitness benefit of inactivating the complex to achieve Bte2 resistance is outweighed by the detrimental impacts on overall physiology.

### Bte2 intoxicates target cells independently of electron flow through the Rnf complex

The dependence of Bte2 on the Rnf complex for target cell intoxication has several potential explanations. One possibility is that Bte2 toxicity is elicited by co-opting the complex in a manner that poisons cellular metabolism. Alternatively, the Rnf complex could serve as a receptor that mediates Bte2 translocation or facilitates its interaction with its target. To distinguish between these possibilities, we sought to determine whether electron flow through Rnf is required for Bte2 intoxication. Using AlphaFold, we obtained a structural model for the Rnf complex of *B. theta* (Fig. 3A). We then identified residues with predicted proximal or direct involvement in electron transfer from reduced ferredoxin and in NAD^+^ reduction by aligning the predicted structure to the solved structure of the *Clostridium tetanomophum* Rnf complex (46) (Fig. 3B and S3B). Substitutions in either iron sulfur cluster-coordinating cysteines in RnfB (necessary for ferredoxin oxidation) or residues in RnfC necessary for NAD+ binding abrogated Rnf function, as measured by growth of mutant strains on Rnf-selective media; however, these mutants remained fully susceptible to Bte2 intoxication (Fig. 3C,D). These data demonstrate that electron flow through the Rnf complex is not required for Bte2 activity.

**Figure 3.**
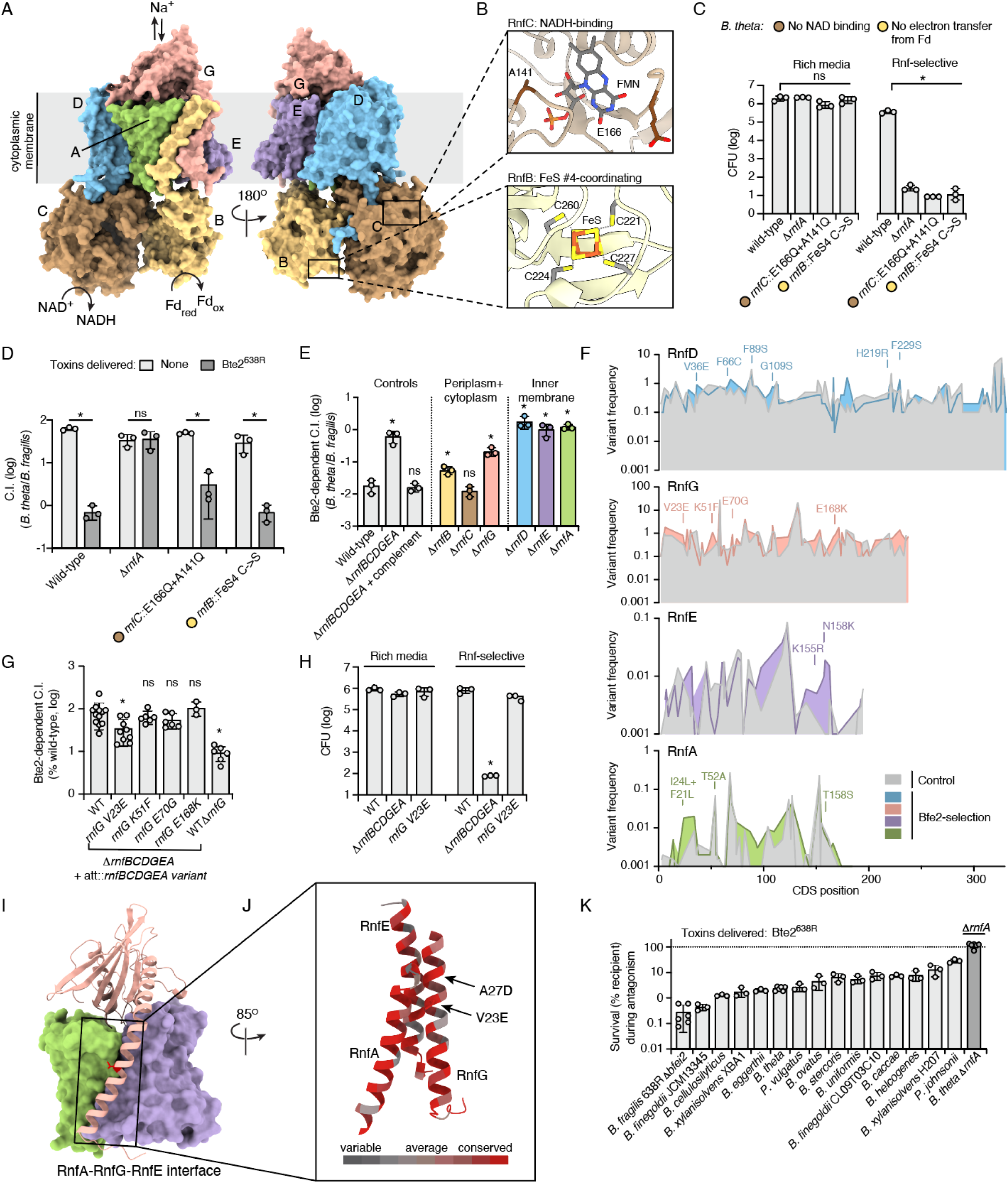
Bte2 reliance on Rnf is independent of electron flow through complex. **A,B)** Alphafold3 multimer prediction for the assembled of *Bt* Rnf complex (RnfBCDGEA). Cofactor binding sites with residues mutated for experiments in (C-D) are indicated in insets (B), overlayed with the Rnf complex structure from *Clostridium tetanomorphum* (PDB: 7zc6) (46). **C)** Growth yields of the indicated strains after propagation on rich (left) or Rnf-selective (right, DMM with propionate and galacturonic acid) media. **D)** Outcome of growth competition assays between the indicated *Bt* strains and *Bf* 638R 11*bte3* (delivers Bte2) or 11*tssC* (no toxins delivered). C.I., competitive index. **E**) Relative fitness of the indicated strains during growth competition assays with *Bf* delivering Bte2^638R^ relative to competition with *Bf* delivering no toxins. **F)** Alternate amino acid frequencies (%) per coding position in the indicated *Bt* Rnf proteins after selection for Rnf function and exposure to Bte2 or control populations selected for only Rnf function, see methods. Mutations enriched by Bte2 exposure and tested for Bte2 resistance in subsequent experiments are indicated. **G)** Relative competitiveness of reconstructed, screen-identified *Bt rnfG* variants during growth competition assays with *Bf* delivering Bte2^638R^ relative to competition with *Bf* delivering no toxins (shown as % WT *Bt* C.I.) (n=3-12). **H**) Growth yields of the indicated *Bt* strains on rich and Rnf-selective media. **I)** Alphafold 3 model of the interface between RnfA (green), RnfG (pink, ribbons) and RnfE (purple) of *Bt.* Residue V23 of RnfG indicated in red. **J)** Zoom in of the interaction between an RnfG helix and RnfA and RnfE, showing the relative conservation of individual residues across Bacteroidales homologs. Location of amino acid substitutions discussed in the text are indicated. **K)** Competitiveness of diverse *Bacteroides* during growth competition assays with *Bf* Bte2^638R^ relative to *Bf* delivering no toxins (%) (n=3-6). Data represent mean ± SD (*p<0.05, unpaired two-tailed Student’s *t*-test).

We next tested whether the proteins constituting the Rnf complex—rather than the catalytic function of the assembly per se—are required for Bte2 toxicity. Owing to T6SS effector delivery to the periplasm, we posited that if Rnf plays a role in Bte2 accessing its target, this would minimally require the inner membrane embedded and periplasmic subunits of the complex. We tested this hypothesis by inactivating each gene in the *rnf* locus of *B. theta* by in-frame deletion and evaluating the susceptibility of these mutants to Bte2. Consistent with a role for Rnf in mediating target access, inactivation of the membrane-embedded subunits of Rnf conferred Bte2 resistance. Interestingly, mutants lacking the largely cytoplasmic subunits, RnfB and RnfC, remained highly susceptible to Bte2. Inactivation of RnfG, the most periplasmically exposed subunit, resulted in an intermediate phenotype.

### Functional Rnf mutants conferring Bte2 resistance are rare

Our findings that Bte2 activity does not require electron flow through Rnf, nor does it require components of the complex localized in the cytoplasm, led us to hypothesize that interaction between Bte2 and membrane-embedded Rnf subunits mediates toxin access to its target. Biochemical experiments aimed at demonstrating a direct interaction between Bte2 and the Rnf complex were unsuccessful; therefore, we performed a genetic selection for Rnf subunit point mutants that confer resistance to Bte2. Focusing on the subunits required for Bte2 intoxication, we employed low fidelity PCR to generate a library containing an average of 3 mutations per clone in a *Bacteroides* integrative vector downstream of the native promoter and genes encoding wild-type copies of the remaining complex subunits (2.7×10^6^ clones, estimated 1.7×10^6^ lacking nonsense mutations) (Fig. S3C). This library was then introduced at an ectopic location of the chromosome in *B. theta* Δ*rnfABCDEFG*. To eliminate clones lacking Rnf function, we propagated the resulting library on Rnf-selective media prior to initiating the selection for Bte2-resistance (Fig. S3D). Only 7.3% of clones survived this initial selection, suggesting that most mutations in Rnf genes are inactivating.

To identify Bte2-resistant (Bte2^R^) variants of the Rnf complex, we passaged the library of RnfDGEA mutants through two rounds of Bte2-specific antagonism by *B. fragilis* 638R 11*bte3* (Fig. S3D). As a control, we passaged it in parallel through two rounds of growth in competition with *B. fragilis* 638R Δ*tssC*. Both pools were subjected to a final round of growth on Rnf-selective media prior to sequencing via a CRISPR-targeted nanopore sequencing-based approach. Comparison of the frequency of amino acid substitution at each position between Bte2-selected and control libraries revealed only a small number of differentially enriched mutations, all of which were rare (<5% frequency) (Figure 3F, Table S2). The most frequent variants we observed were largely detected in *rnfD* and *rnfG*, and at similar frequencies in both the control and Bte2-exposed samples, suggesting they represent mutations that increase fitness relative to the majority of clones in the population rather than confer Bte2-specific resistance (Fig. 3F).

Despite the lack of clear selection for specific Bte2 resistance in our experiment, we proceeded to generate integrative *rnf* complementation constructs bearing mutations enriched following Bte2 selection (Figure S3F). Of the 19 mutants tested, only one, *B. theta rnfG*^V23E^ (detected only after selection with Bte2 and not in control populations), individually conferred significant resistance to Bte2 (Fig. S3E, Fig 3G). Consistent with our selection methodology, this mutation does not impact growth in Rnf-selective media (Fig. 3H). RnfG V23 is predicted to localize to the transmembrane helix anchoring RnfG to the membrane (Fig. 5I-J). Notably, we identified a second Bte2-enriched substitution (A27D, 15-fold) on the same face of the RnfG anchoring helix, one helical turn C-terminal of V23. This region may include an interface exploited by Bte2; however, the limited number of substitutions within the region and its high degree of conservation across species suggests its recalcitrance to functional substitutions (Fig. 3J). Our inability to select variants that both support Rnf-dependent growth and confer robust resistance to Bte2 from an initial collection of over 10^5^ functional clones supports a model in which Bte2 exploits a structurally conserved, functionally essential feature of the Rnf complex to gain access to its target. This is in-line with the high degree of conservation among the genes across *Bacteroides* and closely related genera (90.1% average pairwise identity).To further explore this, we evaluated the Bte2 sensitivity of 15 gut-inhabiting species spanning a wide phylogenetic cross section of the Bacteroidales (Fig. 3K). Though we found a range of susceptibility among these strains, all exhibited significant Bte2-dependent inhibition. When taken together with the results of our genetic screen and our finding that the Rnf complex is broadly required for Bte2 intoxication, this result underscores the exploitability of the Rnf complex for facilitating Bte2-based intoxication.

### Bte2 is a pore-forming effector in the VasX toxin family

Our finding that Bte2 requires the membrane-embedded components of the Rnf complex to exert toxicity suggests its molecular target is either membrane associated or present in the cytosol. To distinguish between these possibilities, we sought to determine the cellular compart in which Bte2 acts. T6SS effectors are delivered to the periplasm of contacting kin cells, where they either exert toxicity directly or translocate to the cytoplasm. Therefore, immunity proteins that prevent self-intoxication from cytoplasmic effectors are essential for survival regardless of T6SS functionality, whereas those associated with periplasmically active effectors are rendered dispensable by T6SS inactivation. We found that *bti2* is readily inactivated in *B. fragilis* 11*tssC*, but not in wild-type cells, indicating that Bte2 acts from the periplasm (Fig. 4A). Furthermore, *bti2* was not essential in the *B. fragilis* 11*rnfA* background, suggesting the Rnf complex mediates access of Bte2 to its target via the periplasm.

**Figure 4.**
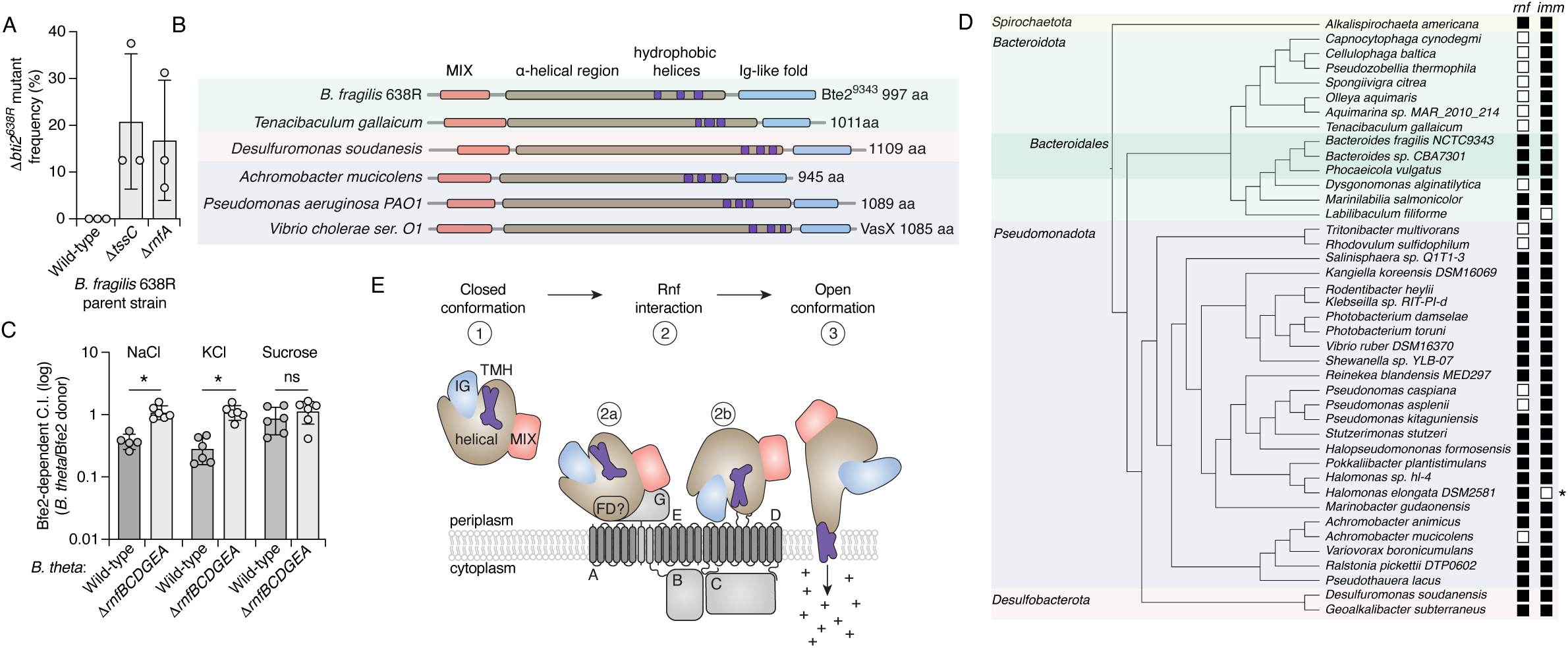
Bte2 is a member of the VasX family of pore-forming toxins. **A)** Frequency of Δ*bti2* generation in the indicated *Bf* 638R backgrounds via 2-step allelic exchange (% vector resolution to Δ*bti2* vs to WT). **B)** Schematic depicted domain architecture of *Bf* 638R Bte2 and VasX-like proteins from diverse phyla identified by Foldseek. **C)** Relative competitiveness of *Bt* (wild-type or Δ*rnfBCDGEA*) during growth competition assays with *Bf* delivering Bte2^638R^ compared to competition with *Bf* delivering no toxins, on media supplemented with different osmoprotectants. **D)** Dendogram of representative species encoding VasX-like toxins. Shading demarks phyla to which species belong. The presence of an Rnf locus and Bte2-immunity protein (*bti2*) within the genome of toxin-encoding strains are indicated. *Indicates a strain containing a degraded T6SS system. **E**) Model of Bte2 intoxication of *Bacteroidales*: 1) Bte2 is delivered into the periplasm in a closed conformation. 2) Bte2 interacts with the Rnf complex, possibly via a flexible domain (‘FD’) in the helical region of Bte2 (a) or the Ig domain (b). The depicted interaction location with respect to Rnf is arbitrary. 3) Bte2 shifts to an open conformation with exposed transmembrane helices (TMH) and the TMH insert into the cytoplasmic membrane to form an ion permeable pore. Data in (A) and (C) represent mean ± SD (*p<0.05, unpaired two-tailed Student’s *t*-test, n=3).

Bte2 family members show little sequence homology to characterized proteins. We thus turned to structural prediction to gain insight into their potential mode of action. AlphaFold modeling followed by Foldseek-based structural similarity searching revealed that Bte2 shares domain architecture and overall structural similarity with members of the VasX toxin family of T6SS effectors (Figure 4B, S4A). Like Proteobacterial VasX family proteins, Bte2 orthologs contain an N-terminal MIX domain for recognition by the T6SS, followed by a large helical region containing predicted transmembrane segments and a C-terminal Ig-like domain (Figure 4B) (18, 47). We speculate that previous studies employing sequence-based approaches to identify VasX family members likely failed to recognize this similarity due to the low primary sequence homology between the Proteobacterial and Bacteroidales proteins (13, 18, 48, 49).

VasX-like effectors form pores that specifically disrupt the electrical component (114¢) of membrane potential by catalyzing monovalent ion influx (18, 50). Accordingly, the toxicity of these proteins is influenced by exogenous ion concentration (18). We measured the ion dependence of Bte2 activity during growth competition experiments between *B. fragilis* and *B. theta*. Similar to the VasX-like toxin Ptx2, Bte2 delivery inhibited competitor growth on media supplemented with NaCl or KCl, and it was ineffective during growth on media containing equimolar sucrose as the primary osmolyte (Fig. 4C) (18).

Our structural and phenotypic data placing Bte2 in the VasX family prompted us to ask whether its members present in species outside of the Bacteroidales also exploit the Rnf complex. We showed that in the case of the Rnf complex-utilizing VasX family member Bte2^9343^, its immunity protein is dispensable in the absence of the Rnf complex. Motivated by this observation, we sought to understand the distribution of *rnf* and cognate immunity genes among strains encoding VasX family members. Given the sequence heterogeneity among these toxins, we used Foldseek to search for proteins sharing structural similarity and domain architecture with Bte2. This led to identification of ∼90 sequence divergent orthologs of Bte2 from four bacterial phyla (Table S4). Analysis of the genomes encoding these proteins identified numerous strains outside of gut-associated Bacteroidales that lack *rnf* genes and nevertheless encode VasX toxins (Fig. 4D, Table S4). This suggests that either these strains do not undergo intercellular self-intoxication via their T6SSs or that the Rnf complex is not required for the activity of these VasX family members. To distinguish between these possibilities, we searched the strains for genes encoding cognate immunity proteins – an evolutionary signature of intercellular self-intoxication for periplasmically acting effectors (3). In strains lacking Rnf, we invariably identified a strongly predicted cognate immunity factor adjacent to each *vasX* gene (Fig 4E). Together, these results suggest that exploitation of the Rnf complex is not a universal feature of VasX family toxins. Instead, we suggest that the distinct metabolic requirements of the gut, including growth on dietary fibers, renders the Rnf complex functionally inescapable and thereby a reliable factor for exploitation.

Taken together with recently published structural analyses of two VasX family proteins, our findings are consistent with a model in which the periplasmic subunits of the Rnf complex are exploited as a receptor that facilitates Bte2 membrane insertion and subsequent pore formation (Fig. 4E). Cryo-EM structures of Ptx2 and Tke5, VasX family proteins deriving from *P. aeruginosa* and *P. putida*, respectively, revealed that their transmembrane helices are enclosed within large α-helical domains in the secreted, soluble forms of the toxins. However, in both cases, authors of the studies noted that structural models predict the transmembrane helices protrude from the protein surface, suggesting significant conformational rearrangement is required to enable membrane insertion (Fig. S4A-C). In support of Bte2 undergoing a similar conformational change upon membrane insertion, AlphaFold2 predicts that the transmembrane helices of Bte2^9343^ are extruded, whereas the algorithm predicts they are enclosed within the α-helical domain in the related protein from *B. fragilis* DS-71 (Fig. S4D). This phenomenon of dual conformations is observed widely among VasX family members (Fig. S4E).

The studies of Ptx2 and Tke5 present different models for how the toxins rearrange to enable membrane insertion. Whitney and colleagues suggest a flexible, α-helical bundle within the larger helical domain may serve as a membrane sensor that triggers the conformational change; Albesa-Jove and colleagues posit the C-terminal, β-sheet-enriched Ig domain fulfills this role (18, 19). We propose that interaction between Bte2 and Rnf, possibly via one of these candidate sensing domains, triggers a conformational change that facilitates Bte2 membrane insertion (Fig. 4E). Perhaps not surprisingly, our attempts to detect interaction between Bte2 and Rnf biochemically were unsuccessful, suggesting the interaction is transient in nature.

## Discussion

Species from the Bacteroidales order live nearly exclusively in the mammalian gut, and thus display numerous adaptations reflecting this specialization. Consistent with the intense interbacterial competition found in this densely colonized habitat, the contact-dependent antibacterial T6SS is prevalent among Bacteroidales (8, 11). However, unlike the broad spectrum targeting displayed by the T6SS^i^ of Proteobacteria, previous studies suggest the Bacteroidales T6SS^iii^ primarily mediates antagonism with other gut Bacteroidales species, their most likely competitors in the gut (13, 22, 51). Our findings suggest this specificity stems at least in part from dependence of the widespread T6SS^iii^ toxin Bte2 on the Rnf complex in target organisms. This complex is conserved in gut-inhabiting Bacteroidales, critical for *in vivo* fitness, and intolerant of mutations, all features that could promote its evolution toward a *Bacteroidales-* targeting specificity determinant. While the Rnf complex is also present in numerous gut Proteobacterial species, their Rnf proteins are sufficiently divergent from those in Bacteroidales to likely preclude Bte2 interaction, and we and others have not observed Bte2-dependent fitness of *Bacteroides* in competition with these organisms (22, 45). Interestingly, a second Bacteroidales T6SS toxin, Bte1, similarly requires a protein target that diverges between Bacteroidales and Proteobacteria, the periplasmic PpiD-YfgM chaperon complex (14). Together, these observations suggest that the mammalian gut environment selects for *Bacteroidales* T6SS toxins specifically effective against *Bacteroidales* targets.

Although the requirement for a receptor in target organisms is rare among T6SS toxins, it is a common feature of narrow-spectrum antibacterial toxins including bacteroicins, the toxins delivered by contact dependent inhibition (CDI), and other secreted toxigenic proteins (52-54). The question thus arises as to what selective pressures drive the targeting spectrum of interbacterial antagonism pathways. In the gut, the range of target specificity exhibited by *Bacteroidales* T6SS toxins may simply reflect the strength of selection imposed by competition between *Bacteroidales* species. By eliminating neighbors that provide benefits, toxins with a broader targeting range may also be maladaptive in this environment. Several studies indicate Proteobacteria can serve as a sink for organic acids and electron donors (hydrogen, succinate) generated during Bacteroidales fermentation of dietary or mucosal polysaccharides, suggesting a benefit to evolving T6SS effectors that avoid collateral damage to this group of organisms (55, 56).

Despite the widespread conservation of the Rnf complex among gut *Bacteroidales*, its physiological role in these organisms is poorly understood. Genomic and metabolomic analyses of *Prevotella* species suggest that in these species, Rnf is critical for reoxidizing reduced ferredoxin during fermentation of glucose to propionate (27). However, this appears to not be the case in *Bacteroides* species, as we show that Rnf is not required for growth on glucose in *B. theta*, nor did addition of exogenous organic acids to shift fermentation toward propionate affect the importance of Rnf. We hypothesize that a recently discovered group B [FeFe] hydrogenase prevalent and highly expressed in *Bacteroides*, but missing from *Prevotella*, is likely responsible for regenerating reoxidized ferredoxin in these organisms (56). Unlike Rnf, this hydrogenase directs the excess electrons from oxidizing ferredoxin to hydrogen gas, rather than using them for NAD^+^ reduction. This is consistent with our finding that Rnf is required for growth, and presumably redox balancing, specifically under conditions that lead to NAD^+^ accumulation. Underscoring the physiological importance of the Rnf pathway, we find it is critical for *B. theta* fitness in mice fed a standard chow diet rich in plant polysaccharides. Future studies of the importance of Rnf in mice maintained on different diets or on a broader range of substrates *in vitro* may help shed light on the specific metabolic pathways employed by *Bacteroidales* during growth on different substrates.

In our model, the Rnf complex mediates Bte2 activity by facilitating conformational rearrangement required for membrane insertion of the toxin transmembrane helices (Fig. 4E). However, our bioinformatic analyses suggest that VasX-like toxins found outside the Bacteroidales do not require the Rnf complex for target cell intoxication, despite these proteins likely also requiring conformational rearrangement to facilitate membrane insertion (18, 19). It is conceivable that VasX family members from Proteobacteria utilize a distinct dedicated receptor or accessory factor. However, the ability of *V. cholerae* VasX to intoxicate eukaryotic cells argues against a strict requirement for a species- or even phylum-specific receptor (49). Interestingly, a recent study of CDI ionophore toxins found these proteins contain variable receptor binding domains that exploit different inner membrane proteins to facilitate membrane insertion of a conserved alpha helical toxin domain (57). The much higher sequence and structural variability among VasX-like toxins precluded a similar categorization of receptor binding domains among these proteins. Further structural characterization of diverse VasX-like toxins and analysis of their delivery requirements will be needed to understand the basis for their targeting specificity across diverse bacteria.

## Methods

### Bacterial strains and culture conditions

All strains used in the study can be found in Table S4. *E. coli* EC100D pir+ (Lucigen) was used for cloning and plasmid amplification, and S17-1 λpir was used for plasmid transfer. *E. coli* strains were grown on lysogeny broth (LB) or LB agar aerobically at 37°C. Unless otherwise noted, *Bacteroides* strains were cultured at 37° C under anaerobic conditions on brain heart infusion agar (BHI; Becton Dickinson) supplemented with 50 μg/ml hemin, 1 μg/ml cysteine HCL, 5% defibrinated horse blood, and 0.2% sodium bicarbonate (referred to as rich media in the text) (58). BHI broth supplemented with 0.2% sodium bicarbonate and 1 μg/ml cysteine (BHIS) was used as liquid media for culturing and normalizing strains unless otherwise noted. Anaerobic culturing was performed either in an anaerobic chamber (Coy Laboratory Products) filled with carbon dioxide 10%, hydrogen 10%, nitrogen 80% (Praxair), or in Becton Dickson BBL EZ GasPak chambers. Antibiotics and chemicals were added to media as needed at the following concentrations: carbenicillin 150 μg/mL (*E. coli*), kanamycin 50 μg/ml (*E. coli*), gentamicin 60 or 200 μg/mL^−1^ (standard *Bacteroides* growth media and media for conjugations, respectively), gentamicin 60 μg/mL (*E. coli*/*Bacteroides*), erythromycin 12-25 μg/mL (*Bacteroides*), tetracycline 6 μg/mL (*Bacteroides*), floxuridine (FUdR) 200 μg/mL (*Bacteroides*). Varel-Bryant medium (58) without glucose was used as the DMM for *Bacteroides* and supplemented with the following as needed: D-glucose (5mg/ml), D-galacturonic acid (5 mg/ml), L-Rhamnose (5 mg/ml), D-fructose (5 mg/ml), sodium propionate (1 and 5 mg/ml, for screening and growth assays, respectively), lactate (5mg/ml), formate (5mg/ml), acetate (5mg/mL), and succinate (5mg/mL). DMM supplemented with propionate and galacturonic acid is referred to as Rnf-selective media.

### Plasmid Construction

Plasmids and oligonucleotides used in this study can be found in Table S4. Plasmids were designed using Geneious Prime and all primers were synthesized by Integrated DNA Technologies (IDT). Plasmids were constructed using Gibson assembly and standard molecular procedures, and confirmed by sequencing. Plasmids were transformed and maintained in *E. coli* EC100D under carbenicillin or kanamycin selection. Constructs for introducing deletions and point mutations in the chromosome of *Bacteroides* strains were generated using 0.5-1 kb regions of homology flanking the modification site in the *Bacteroides* chromosome and assembled into the pExchange vector using Gibson assembly, as previously described (51, 59, 60). For strain marking and complementation, the pNBU2 integrative vector, which integrates at the *att* sites, was assembled with the sequence of interest under the native promoter (61, 62).

### Genetic manipulation of *Bacteroides* strains

To generate genetically modified *Bacteroides* strains, vectors were mobilized from *E. coli* S17-1 into *Bacteroides* strains via conjugation in aerobic (for gene deletion) or anaerobic (for marking and complementation) conditions. To improve the frequency of plasmid introduction into *Bacteroides*, *E. coli* S17-1 expressing RPK31 (63) was included in the conjugations at a 1:1:1 ratio with *E. coli* S17-1 containing the vector of interest and the recipient *Bacteroides* strain. Each of these strains was normalized to 150 ul of OD 600 = 3, pooled and plated onto rich media without antibiotic supplementation. After overnight conjugation, *Bacteroides* clones with the integrated vectors were isolated on rich media with erythromycin or kanamycin selection. For gene deletion, two-step allelic exchange with the pExchange vector was utilized (60). Following mobilization into *Bacteroides*, pExchange merodiploids were isolated on rich media with erythromycin selection and further passaged on FudR counter-selection to resolve the thymidine kinase (*tdk*) containing pExchange backbone. *Bacteroides* mutants were confirmed by colony PCR followed by Sanger sequencing.

### Interbacterial competition assays

Strains to be employed for these assays were struck onto solid media from single colonies. Following overnight growth, colonies of the antagonizing (donor) and target (recipient) strains were scraped from the plate and resuspended in liquid media to OD_600_ = 100 (donor) or 1 (recipient). Equal volumes of the two suspensions were then combined and 10 μl samples of the mixture were spotted on 3% (w/v) agar rich media with gentamycin (60 mg/mL) for standard assays. In osmolyte addition assays, mixed suspensions were spotted on BHI agar without dextrose supplemented with 300 mM NaCl, sucrose, or KCl. Competitions were incubated anaerobically for ∼18-22 h at 37°C as previously described (8, 45, 51, 59), and aliquots of the mixed strain suspensions were diluted and plated on media selective for the donor and recipient populations (erythromycin or tetracycline added) to enumerate initial colony forming units (CFU).

After incubation, competitions were scraped from solid media and resuspended in BHIS liquid media, serially diluted, and plated on rich media containing erythromycin or tetracycline selection to separate and enumerate the CFU of each strain after competition. The competitiveness of donor and recipient strains was determined by calculating the competitive index (C.I., CFU_f_ recipient/CFU_i_ recipient)/(CFU_f_ donor/ CFU_i_ donor). Toxin delivery-dependent competitiveness of recipient strains was calculated as the C.I. following competition with an antagonizing strain relative to the C.I. with a T6SS-inactivated strain (Δ*tssC*). All competitive growth experiments were performed with at least biological triplicates and at least two independently replicated experiments.

### RB-Tn-Seq screen and analysis

To initiate the RB-Tn-Seq screen, we grew overnight cultures of a *B. theta* barcoded transposon library containing 315,668 uniquely barcoded mutants (20) and T6SS variants of *B. fragilis* 638R and *B. fragilis* NCTC9343. Overnight cultures were spun down an resuspended in fresh medium to OD_600_ = 100 (*Bf* donor strains) or 1 (*Bt* library). Equal volumes of individual donor strains were mixed with the library, and 50 replicate spots (10 μl) of each pair was plated on 3% (w/v) agar rich media with gentamycin (60 mg/ml). This allowed for a library population size at least 1000-fold greater than the number of unique clones at the initiation of the experiment, and at least 20 fold greater under maximal intoxication. Competitions were incubated anaerobically at 37C for 20 hours before scraping the competitions from the plates and resuspending in liquid media. Aliquots of these suspensions were plated diluted and plated on rich media supplemented with erythromycin (*Bt*) or tetracycline (*Bf*) to enumerate surviving cells, and genomic DNA was extracted from the remainder of the suspensions (Qiagen DNeasy Blood & Tissue Kit). The library barcodes were then amplified (Q5 High-Fidelity DNA polymerase, NEB) as previously described, utilizing TruSeq indexes for automatic demultiplexing of the samples by Illumina ((21),primer set 2). Amplified barcodes were purified (Zymo DNA Clean and Concentrator Kit), quantified (Qubit), and libraries were pooled and sequenced on an Illumina MiSeq V3-150 flow cell as single-end reads with 2% phiX DNA spiked in. Illumina sequencing reads were analyzed using a custom Python script and the Fitness Browser resources (https://morgannprice.org/FEBA/Btheta/). Briefly, reads per barcode were tallied for each sample and the sum of the reads for all of the insertions per coding sequence were recorded. Insertions per gene were normalized by gene length and median insertion abundance/sample.

### Identification of Bte2 orthologs

To identify Bte2 orthologs in Bacteroidales (taxID: 171549), psiBLAST on Bte2^638R^ was run to convergence with the NCBI nr database and filtered for 90% query coverage (E-value <0.005). A tree was generated using the NCBI distance tree feature, with a midpoint root, and leaves collapsed as indicated (Figure S1G).

### Monosaccharide and fermentation end-product supplementation growth assays

*Bt* strains were grown on solid rich media overnight from single colonies and normalized in phosphate buffered saline (PBS) to 10^9^ CFU/ml. Strains were further diluted prior to plating such that ∼2000 CFU were plated onto the test DMM with supplements. An aliquot of the starting suspension was separately titered on rich media to determine the input colony forming units (CFU). Following anaerobic incubation, colonies were harvested by flooding the plates with 2 x 1mL PBS, scraping the colonies and resuspending in a separate tube. Viable CFU/plate was determined by serial dilution onto a non-selective rich media, and normalized by the input CFU for each strain.

### Gnotobiotic mouse experiments

#### Strain preparation and mouse colonization

Germ-free 6–12 week-old male and female Swiss Webster mice from multiple litters were randomized, housed in pairs in single Techniplast cages with a 12-hour light/dark cycle, and fed a standard lab diet (Laboratory Autoclavable Rodent Diet 5010, LabDiet) ad libitum, in accordance with guidelines approved by the University of Washington Institutional Animal Care and Use Committee. The number of animals were selected based on Lemorte power calculations to measure a significance difference of ∼10-fold bacterial abundance with a *p* < 0.05 at 80% power along with housing and maintenance considerations for gnotobiotic mice. Mice were confirmed to be sterile prior to colonization by PCR with primers targeting the 16S rRNA gene.

Germ-free mice were colonized by oral gavage with bacteria grown on rich media plates supplemented with gentamycin (60mg/ml) or glycerol stocks of a wild mouse microbiome supplied by Taconic (WildR) (42). To prepare *B. fragilis* strains for gavage, single colonies were restruck on plates and incubated anaerobically at 37 °C for 24 hours. Cells were harvested and diluted to 10^9^ CFU/ml in pre-reduced BHI broth. Mice colonized with two bacterial strains were gavaged with a total of 5×10^8^ CFU/200 µL at a 10:1 ratio of *B. fragilis* wild-type or Δ*tssC* to *B. fragilis* Δ*ei2*. To colonize mice with *B. fragilis* and the wild mouse microbiome, *B. fragilis* strains were mixed at a 1:1 ratio (either *Bf* WT with *Bf* Δ*ei2,* or *Bf* WT with *Bf* Δ*rnfA)* and diluted to ∼2×10^9^ CFU/mL of each *Bf* strain in 1 mL of the wild mouse microbiome glycerol stock (containing ∼8*10^8^ CFU of the WildR community) under anaerobic conditions. Mice were gavaged with 150µl of the WildR:*Bf*:*Bf* strain mixtures.

#### Quantification of B. fragilis from mice

Fecal pellets were collected over time and upon sacrificing each mouse, cecal contents were also collected and stored at -80°C. For quantification of *B. fragilis*, thawed fecal pellets or cecal contents (∼50-100 mg/sample) were added to tubes containing 200 µL of 0.1-mm-diameter zirconia/silica beads (BioSpec Products, Bartlesville, OK), 500 µL of CP Buffer (Omega Biotek, #PDR042), 250 µL of 20% SDS, and 550 µL of phenol:chloroform:isoamyl alcohol (24:24:1). Microbial cells were mechanically lysed with a bead beater at room temperature (BioSpec Products; instrument on high for 1 min; 2 cycles). Isolated gDNA was column purified using E-Z 96^TM^ DNA plates (Omega Biotek; #BD96-01) following the standard protocol and quantified by fluorimetry (Qubit, ThermoFisher). The relative abundance of each *Bf* strain was monitored in purified fecal DNA from each mouse at each timepoint by qPCR with strain specific primers (Table S4). qPCR was performed using a CFX Connect Real-Time System (Bio-Rad) and SYBR FAST universal master mix (KAPA Biosystems). Strain quantities were calculated using a standard curve of purified genomic DNA.

### Structure modeling and conservation analysis of the Rnf Complex

Alphafold3 was used to predict the structure of the multimeric Rnf complex from *B. theta* (64). To identify conserved residues involved in electron transfer in RnfB (FeS cluster #4) and RnfC (NADH/NAD^+^ binding), this prediction was aligned to the cryoEM structure of the Rnf complex from *C. tetanomorphum* (PDB: 7zc6 (46)) using ChimeraX 10.1.

To determine the conservation at each position in the *B. theta* Rnf complex, the amino acid alignments of the top 250 Bacteroidales (taxID: 171549) hits by NCBI pBLAST were extracted for each subunit. Consurf was used to determine the level of conservation at each residue and to map the conservation scores onto the predicted structures for each subunit (https://consurf.tau.ac.il/consurf_index.php). The subunits were assembled using ChimeraX 10.1 with the multimerized prediction of the *Bt* Rnf complex as a scaffold.

To generate a conservation score for each subunit of the Rnf complex in gut-relevant *Bacteroidales*, the sequences of Rnf complex subunits from strains representing all major families of Bacteroidales that reside primarily in the mammalian gut were extracted. These sequences were aligned in Geneious, the pairwise identity (%) was calculated for each subunit, and the average identity was calculated across subunits to obtain an overally similarity score. A similar analysis was used to generate similarity scores for the Rnf proteins of the 15 strains of Bacteroidales that were demonstrated to be Bte2^638R^-sensitive in this work.

### Genetic screen for functional, Bte2-resistant Rnf complexes

#### Rnf gene mutagenesis and library construction

The genes encoding RnfDGEA were amplified from *B. theta* VPI-5482 gDNA with the Diversify™ PCR Random Mutagenesis Kit (Takara Bio) to introduce random mutations at an average frequency of 3 mutations per *rnfDGEA* sequence. This mutation rate per clone was selected to strike a balance between minimizing the frequency of wild-type *rnfDGEA* sequences in the input library, while avoiding a high rate of nonsense and Rnf inactivating mutations that would decrease the diversity of functional Rnf complexes in the library. Mutagenized sequences were assembled with Gibson Assembly Master Mix (NEB) into the integrative vector pBNBU2_erm, downstream of the wild-type Rnf promoter and *rnfBC,* resulting in a library of pNBU2_erm_P*rnfBC*_mut*DGEA* vectors. The assembled vectors were transformed into TransforMax™ EC100™ Electrocompetent E. coli (Lucigen) to produce a library of ∼2 x 10^6^ assembled vectors. The vector library was then tranformed into *E. coli* S17-1 for mating into *B. theta* 11*tdk* 11*rnfBCDGEA*. ∼2.8 x 10^6^ transconjugants of *B. theta* 11*tdk* 11*rnfBCDGEA att:pNBU2_erm_PrnfBC_mutDGEA* were obtained. The pooled transconjugants were passaged on Rnf-selective media (DMM with with D-galacturonic acid (5mg/ml) and sodium propionate (1mg/ml)) to select for Rnf complexes that have retained Rnf function, resulting in an input library (Library S1) of ∼ 2 x 10^5^ clones to screen for Bte2-resistance.

#### Selection for Bte2-resistance

Interbacterial competition assays were conducted with *B. theta* library S1 (OD_600_ = 1) and either *B. fragilis* 638R Δ*tssC* (no toxin delivered, OD_600_ 100) or *B. fragilis* 638R 11*btei1* (Bte2 delivered, OD_600_ = 100). Competitions were set up as described above for interbacterial competition assays except PBS was used as the liquid media for normalizing cells. Each competition was harvested and plated on rich media (BHIS/blood/erm25) to select for *B. theta*. The resulting library pools were then each passaged through a second round of competition with either *B. fragilis* 638R Δ*tssC* (no toxin delivered) or *B. fragilis* 638R 11*btei1* (Bte2 delivered), and the resulting library pools were harvested on rich media (BHIS/blood/erm25) to select for *B. theta*. At both competition passages, *B. theta* was titered on both rich media and Rnf-selective media, and the number of 10 μl competitions were scaled to maintain a >100x excess of the predicted number of unique *B. theta* clones throughout the competition. The resulting libraries were then passaged on Rnf-selective media to remove Rnf complexes with inactivated function from the library, to produce the final libraries. After each passage and selection, individual clones were sequenced with sanger sequencing for quality control (Azenta). DNA from each of the pooled libraries after passaging and selection was extracted in preparation for sequencing (Qiagen Blood and Tissue Kit).

#### Sequencing of mutagenized rnf complexes

For each selection/passage analyzed, targeted long-read sequencing of the region of interest from the integrated vector containing the mutagenized *RnfDGEA* locus was performed using Oxford Nanopore’s Cas9 Sequencing Kit (SQK-CS9109, Oxford Nanopore) (https://community.nanoporetech.com/docs/prepare/library_prep_protocols/cas9-targeted-sequencing/v/enr_9084_v109_revv_04dec2018) using two crRNAs targeting to each side of the *rnfDGEA* sequence (Table S4). The crRNAs, tracrRNA and purified Cas9 were obtained from IDT.

Sequencing was performed using MinION Flow Cells R10.4.1 on a Minion Mk1B sequencer (Oxford Nanopore) according to manufacturer’s instructions, (36 h run time per sample). Basecalling was performed using the MinKNOW software (need version #). Reads were mapped to a reference genome (*B. theta* Δ*tdk* Δ*rnfBCDGEA* att:pNBU2_*rnfBCDGEA*) using MiniMap2 2.24 in Geneious and filtered for alignments covering both ends of the integrated target sequence. Variant calling was performed using Geneious Prime (v 2025.1.1) with subset of 50-60k reads per sample (randomly selected from all reads) with default settings. Frequencies of amino acid substitutions at each position in the coding sequence of mutagenized genes were calculated by summing the frequency of all alternate alleles resulting in amino acid substitutions at that position. The frequency of substitution at each position in clones exposed to Bte2 (two rounds of selection) was compared to the average frequency of substation observed in control samples (input library pool after selection for Rnf function and library pool competed with *B. fragilis* Δ*tssC*).

### Distribution of *rnf* and *bti2* in strains with VasX-like proteins

To assess the extent of correlation between genomes that encode VasX-like toxins and the Rnf complex, proteins with a similar domain architecture to Bte2 were first identified in the AFBD50 using a Foldseek search of the Alphafold3 model of Bte2 from *B. fragilis* 9343 (65). Manual inspection revealed that those orthologs with regions of similarity to Bte2 at least 850 amino acids in length contained the C-terminal Ig domain and were selected as bonafide Bte2 orthologs (93 orthologs identified). Genomes for the organisms containing these candidates were identified by performing a function search for the VasX-like Pfam (pfam20249) in the IMG database, and cross-referencing positive genomes against the orthologs identified in Foldseek, leading to identification of 72 genomes encoding Bte2 orthologs. These genomes were then screened for the presence of two COGs diagnostic of the Rnf complex, COG4657 (RnfA) and COG2878 (RnfB). Of the 72 VasX-encoding genomes examined, we found 20 that lacked *rnf* genes; a representative subset of examined genomes are depicted in Fig. 4D.

### Structural modeling of Bte2-family proteins

Alphafold3 was used to predict the structures of representatives of phylogenetically diverse proteins with shared domain architecture to Bte2, identified by Foldseek (described above). The coordinates of each shared domain was mapped to these orthologs using multiple methods. MIX domains were identified using NCBI conserved domains (cd20705-cd20709). The central α-helical region was identified by eye, as was the distinctive c-termal Ig-like fold that consistently forms an isolated β-sheet structure that is separate from the helical region. The transmembrane helices within the helical domain were mapped using using DeepTMHMM - 1.0 (https://services.healthtech.dtu.dk/services/DeepTMHMM-1.0/) (66).

### Statistics and reproducibility

Statistical tests were performed using Graphpad Prism. For bacterial growth assays and competition assays, the statistical tests used and the number of replicates collected in parallel on a single day are indicated in the corresponding methods or figure legends. Each experiment was also replicated at least once on separate days with at least n=3 per replicate. Statistical methods were not used to predetermine sample size, and randomization and blinding were not employed. Significance of differences between mouse groups at each timepoint were determined using a two-way ANOVA with a multiple comparisons test.

## Acknowledgements

Thank you to Adam Deutschbauer for the barcoded transposon library of *Bacteroides thetaiotaomicron,* David Brinkley, Yaxi Wang, Qinqin Zhao, Pooja Srivinas, and Lucy Brennan for feedback on the manuscript and figures, Frank DiMaio for assistance with structural prediction analysis and Carrie Harwood and members of the Mougous laboratory for helpful discussions. H.K.R. is an HHMI awardee of the Life Sciences Research Foundation. J.D.M. is an HHMI Investigator and holds the Lynn M. and Michael D. Garvey Endowed Chair in Gastroenterology at the University of Washington.

## Figure Legends

**Supplemental Figure 1.**
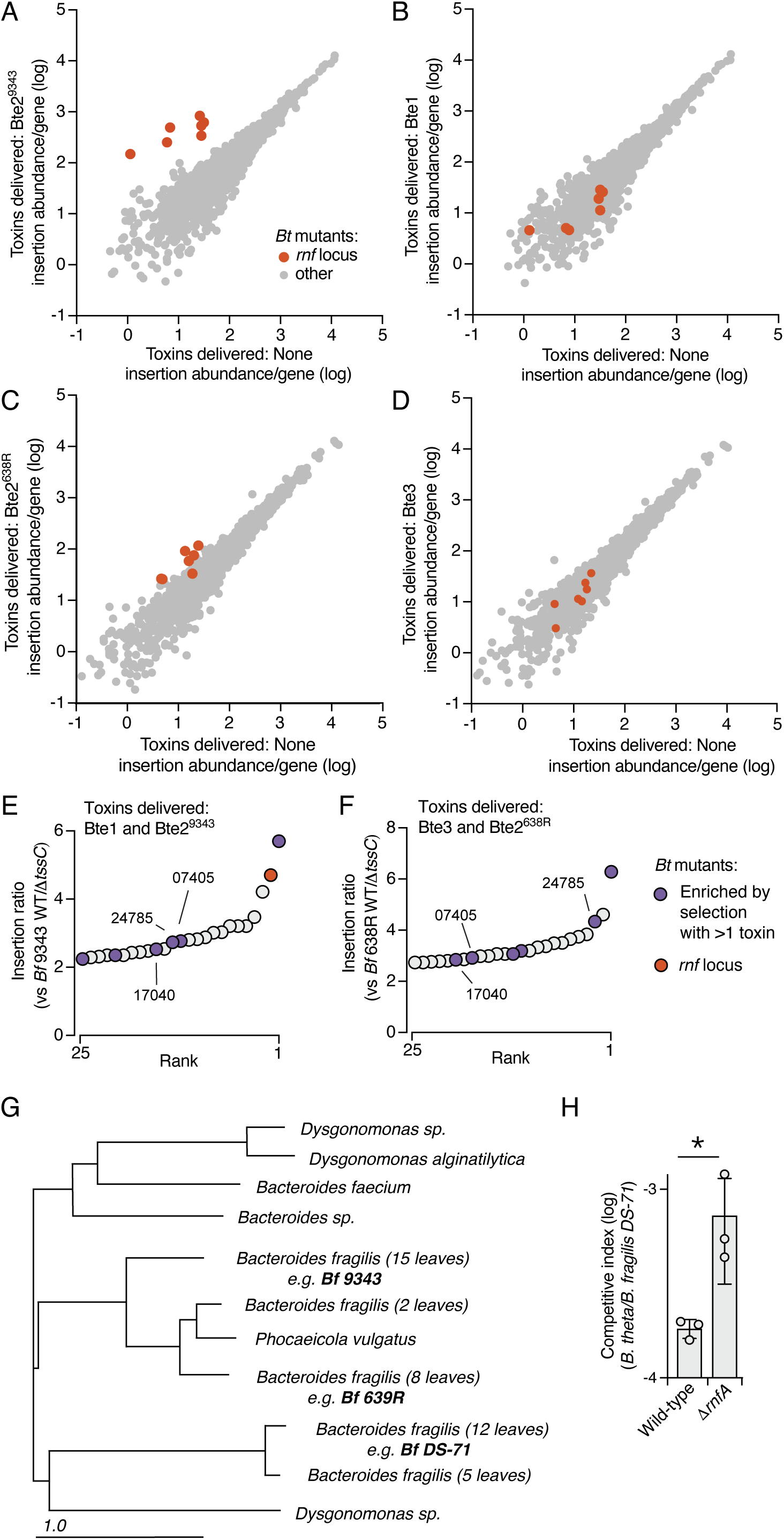
Transposon mutant screening and growth competition assays to identify *Bacteroides* T6SS intoxication susceptibility determinants. **A-D)** Transposon insertion abundance per *Bt* gene during antagonism with *Bf* strains delivering the indicated individual T6SS toxins (y-axis) or *Bf*Δ*tssC* (inactivated T6SS, x-axis). **E-F)** Rank fold enrichment of *B. theta* (*Bt*) transposon insertion mutants following growth in competition with the indicate WT *B. fragilis* strains*. Bt* mutants specifically mentioned in the text are highlighted BT_RS24785, AAA-family ATPase; BT_RS07405, hypothetical; BT_RS17040, predicted glycosyltransferase. **G)** Phylogeny of Bte2 family proteins in *Bacteroidales*. Bte2 orthologs tested for Rnf-dependence via competition growth assays are indicated in bold. **H)** Outcome of growth competition assays between *Bt* (wild-type or Δ*rnfA*) and *B. fragilis* DS-71. Data represent mean ± SD (*p<0.05, unpaired one-tailed Student’s *t*-test, n=3).

**Supplemental Figure 2.**
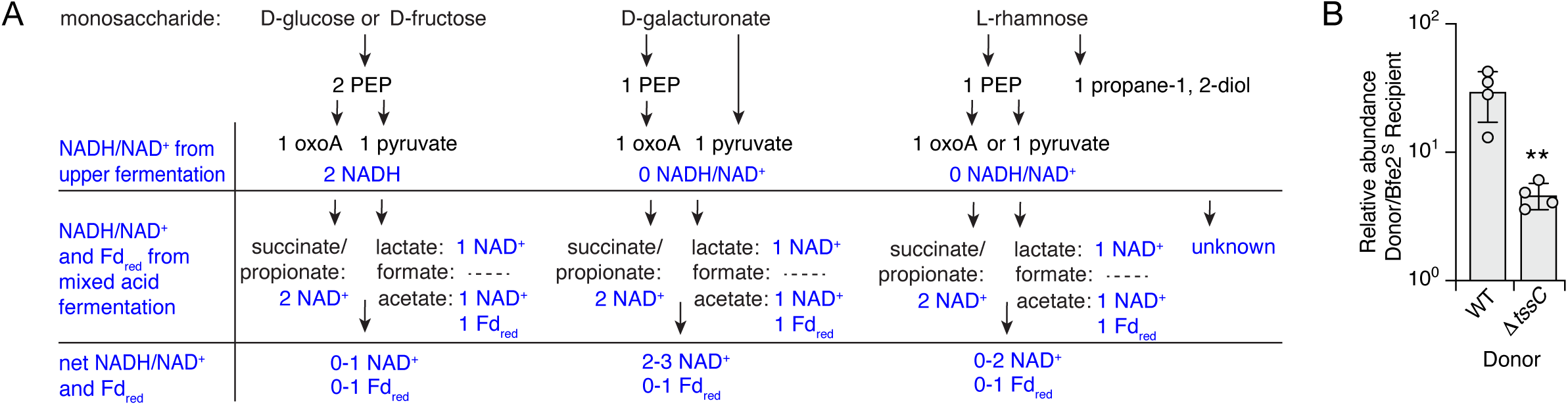
*B. theta* cofactor utilization varies during growth on different monosaccharides and Bte2 improves *Bf* fitness in vivo. A) Predicted NADH/NAD^+^ and Fd_red/ox_ utilization in *Bt* as determined by Kegg pathway analysis during growth on the indicated monosaccharides. B) Abundance of *Bf* (*Bf* 638R WT or Δ*tssC*) relative to a Bte2-susceptible variant *(Bf* 638R Δ*btei2*) in cecal contents 10 days after pairwise gavage into gnotobiotic mice. Data represent mean ± SD (**p<0.01, unpaired two-tailed Student’s *t*-test, n=4).

**Supplemental Figure 3.**
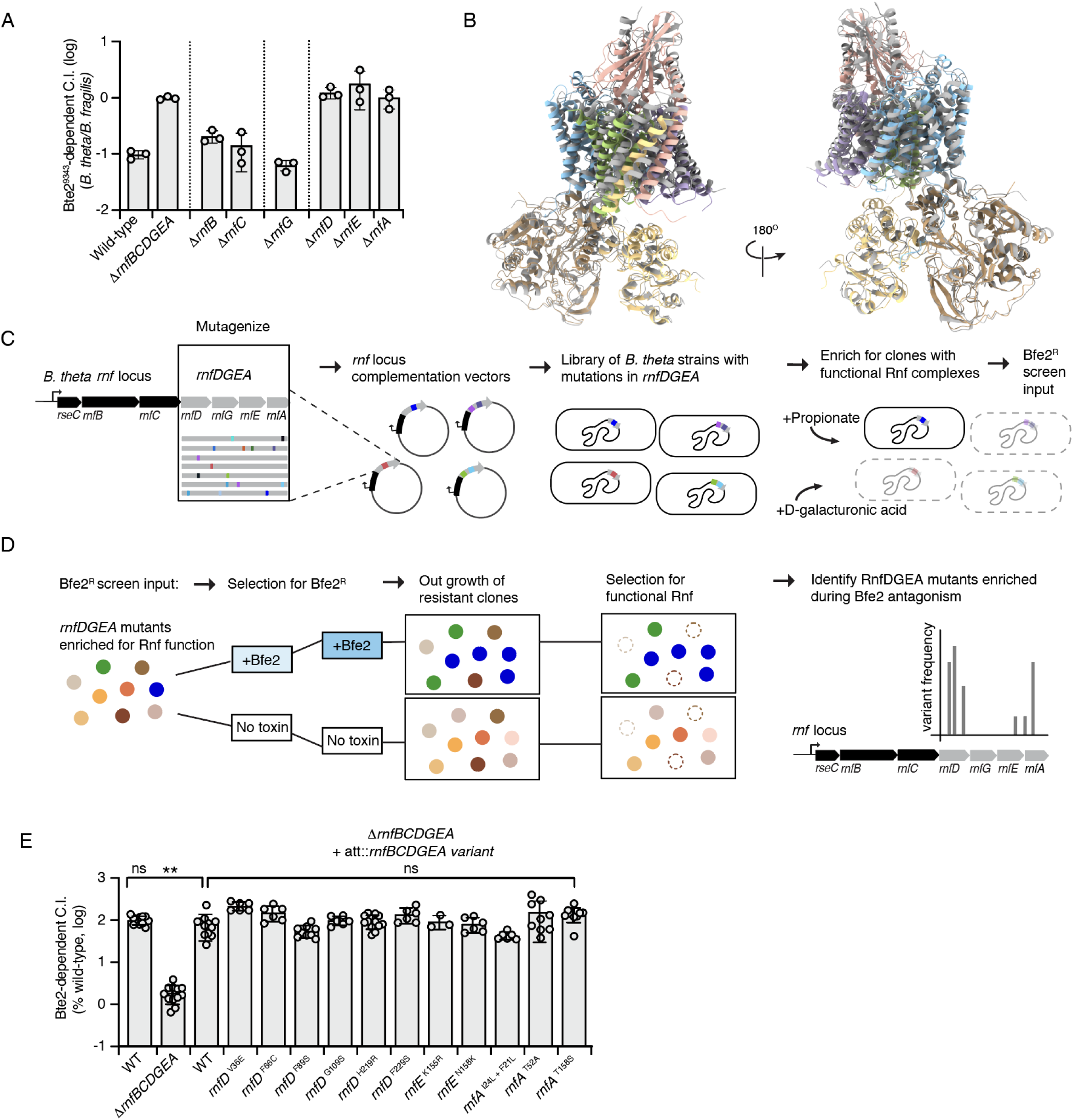
RnfDEA are not permissive to mutations that resist Bte2 and maintain Rnf function. **A)** Competitiveness of *Bt rnf* locus variants during growth competition assays with *Bf* delivering Bte2^9343^ relative to in similar assays with *Bf* delivering no toxins. **B)** Alphafold3 multimer-generated model of the *Bt* Rnf complex (subunits colored according to Figure 2A) overlayed with the solved structure of the Rnf complex from *Clostridium tetanomorphum* (grey, PDB: 7zc6 (46)). **C,D)** Schematic depicting construction of a library of *Bt* with randomly mutagenized *rnfDGEA* subunits and initial selection for Rnf complex function (C) and design for the screen to identify Bte2-resistant variants in the library of *Bt* with mutagenized *rnfDGEA* (D). **E)** Relative competitiveness the indicated reconstructed variants of *Bt rnfDEA* during growth competition assays with *Bf* delivering Bte2^638R^ relative to competition with *Bf* delivering no toxins (shown as % WT *Bt* fitness) (n=3-12). Variants represent those selected by Bte2 exposure during our genetic screen. Data represent mean ± SD (**p<0.05, unpaired two-tailed Student’s *t*-test, n=3).

**Supplemental Figure 4.**
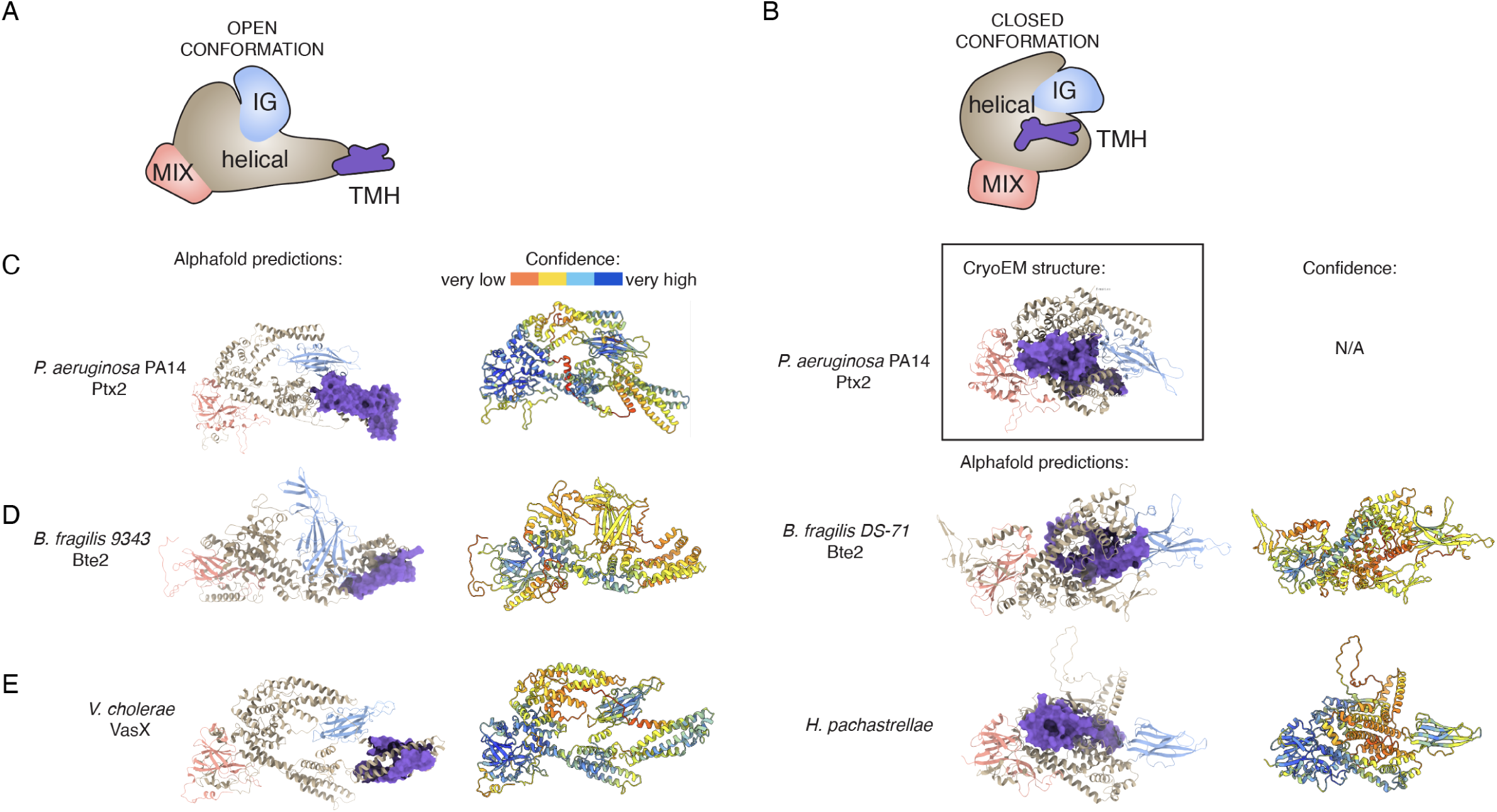
Structure modeling of VasX-family proteins predicts variable conformations for the helical domain. **A, B**) Schematics depicting a generalized VasX-family protein in the open (**A**) or closed (**B**) conformations. **C**) Alphafold3 prediction (left) and CryoEM structure of Ptx2 from *P. aeruginosa* (right). Model predictions are colored by domains (left, colors correspond to A and B) or model confidence (right). **D, E**) Alphafold3 models of representative VasX family proteins, colored by domains and model confidence. Examples were selected to indicate related proteins from *Bacteroides fragilis* (**D)** or γ-Proteobacteria (**E**) that have different predicted conformations.

## Notes

### Competing Interest Statement

The authors have declared no competing interest.

